# Dynamic changes in the mouse hepatic lipidome following warm ischemia reperfusion injury

**DOI:** 10.1101/2022.07.10.499482

**Authors:** Kim H.H. Liss, Muhammad Mousa, Shria Bucha, Andrew Lutkewitte, Jeremy Allegood, L. Ashley Cowart, Brian N. Finck

## Abstract

Liver failure secondary to nonalcoholic fatty liver disease (NAFLD) has become the most common cause for liver transplantation in many parts of the world. Moreover, the prevalence of NAFLD not only increases the demand for liver transplantation, but also limits the supply of suitable donor organs because steatosis predisposes grafts to ischemia-reperfusion injury (IRI). There are currently no pharmacological interventions to limit hepatic IR injury because the mechanisms by which steatosis leads to increased injury are unclear. To identify potential novel mediators of IR injury, we used liquid chromatography and mass spectrometry to assess temporal changes in the hepatic lipidome in steatotic and non-steatotic livers after warm IRI in mice. Our untargeted analyses revealed distinct differences between the steatotic and non-steatotic response to IRI and highlighted dynamic changes in lipid composition with marked changes in glycerolipids and glycerophospholipids. These findings enhance our knowledge of the lipidomic changes that occur following IRI and provide a foundation for future mechanistic studies. A better understanding of the mechanisms underlying such changes will lead to novel therapeutic strategies to combat IR injury.

## Introduction

Obesity-associated nonalcoholic fatty liver disease (NAFLD) is one of the most common causes of chronic liver disease and one of the leading indications for liver transplantation (1, 2). Additionally, as steatotic livers are more susceptible to ischemia reperfusion injury (IRI), the increasing prevalence of NAFLD also limits the supply of donor livers deemed suitable for liver transplantation (3, 4). Ischemia reperfusion injury (IRI) occurs when there is a temporary interruption in organ perfusion followed by re-establishment of blood flow. It is unavoidable in most liver-related surgeries including hepatic resections and liver transplantation. This leads to organ injury and in the case of liver transplantation, can result in primary graft non-function and early allograft dysfunction (5). Due to increased susceptibility to IRI, the use of steatotic livers in liver transplantation has been associated with inferior patient and graft outcomes (4, 6, 7).

However, the mechanisms that lead to increased susceptibility of steatotic livers to IRI is not well understood and there are currently no pharmacological interventions to prevent or treat IRI. Studies in non-steatotic liver have indicated that IRI is associated with alterations in lipid metabolism (8–14). Lipids play important physiological functions as an energy source, as signaling intermediates, and as building blocks for plasma membranes (15, 16). In addition to their crucial role in normal physiological processes, dysregulation of lipid metabolism and alterations in lipid composition have been recognized in pathological states such as metabolic syndrome, cancer pathophysiology, immune dysregulation, inflammatory states, and age-related diseases (17–20). Although alterations in lipid abundance and compositions have been noted after IRI, how the presence of underlying steatosis impacts dynamic changes in the hepatic lipidome is not well defined. Indeed, comprehensive hepatic lipidomic analyses comparing steatotic and non-steatotic responses to IRI have not been sufficiently addressed.

The present study was conducted under the premise that characterization of the dynamic changes in lipid composition and metabolism in steatotic and non-steatotic liver might identify novel pathophysiologic mediators of IRI. We used a well-established mouse model of warm hepatic IRI and performed unbiased, untargeted, comprehensive lipidomic analysis of steatotic and non-steatotic livers exposed to IRI at several time points after reperfusion. The abundance of several lipids changed dramatically after IRI and many of these were also affected by preexisting steatosis. This could facilitate identification of novel biomarkers and mechanistic targets for drug development and therapeutic intervention to ameliorate IRI.

## Materials and Methods

### Animals

Male C57BL/6J mice were purchased from Jackson Laboratory (Bar Harbor, ME). At 6 weeks old, mice were continued on standard chow diet (PicoLab Rodent diet 205053) or transitioned to a diet with high fat (42% calories), sucrose (34% calories), and cholesterol (0.2% w/w) (42% HF; TD 88137, Envigo, Indianapolis, IN). Mice were maintained on diet for 8 weeks prior to surgery. All animal studies were approved by the Institutional Animal Use and Care Committee of Washington University School of Medicine and comply with the *Guide for the Care and Use of Laboratory Animals* as outlined by the National Academy of Sciences.

### Hepatic Ischemia Reperfusion Surgery

Hepatic ischemia was induced using a 70% ischemia model as previously described (Abe 2009; Liss 2021; Liss 2018). Briefly, mice were anesthetized using isoflurane inhalation. Midline laparotomy was performed followed by cross-clamping of the hepatic artery, portal vein, and bile duct distal to the branch point to the right lateral lobe to induce ischemia to the median and left lobes. The atraumatic clamp was released after one hour followed by 6, 24, and 72 h reperfusion. Mice undergoing sham surgery underwent midline laparotomy with vascular clamping and were maintained under isoflurane anesthesia for one hour. At the predetermined reperfusion time point, mice were euthanized and plasma and liver samples were collected for analysis.

### Plasma parameters

Plasma alanine aminotransferase was measured using a commercially available colorimetric kinetic assay (Teco Diagnostics, Anaheim, CA) according to manufacturer’s instructions. Plasma nonesterified fatty acids, total cholesterol, and triglycerides were measured using commercially available colorimetric kits (Wako Diagnostics, Mountain View, CA; and Thermo Fisher Scientific) according to the manufacturer’s instructions.

### Histology

A portion of the left lateral lobe was harvested at the time of sacrifice and placed in 10% neutral buffered formalin followed by 70% ethanol. The tissues were then embedded in paraffin, sectioned, and stained with hematoxylin-eosin (H&E) stain.

### Untargeted lipidomics

Internal standards were purchased from Avanti Polar Lipids (Alabaster, AL) as their premixed SPLASH LIPIDOMIX mass spec standard. Internal standards were added to samples in 10 μl aliquots. Standards included 15:0-18:1(d7) PC, 15:0-18:1(d7) PE, 15:0-18:1 (d7) PS, 15:0-18:1 (d7) PG, 15:0-18:1 (d7) PI, 15:0-18:1 (d7) PA, 18:1 (d7) LPC, 18:1 (d7) LPE, 18:1(d7) cholesterol ester, 18:1(d7) MAG, 15:0-18:1(d7) DAG, 15:0-a8:1(d7)-15:0 TG, 18:1(d9) SM, and cholesterol (d7). For LC-MS/MS analyses, a Thermo Scientific Q Exactive HF Hybrid Quadrupole-Orbitrap Mass Spectrometer was used. Samples were separated via a Thermo Scientific Vanquish Horizons UHPLC System functioning in binary mode.

Samples were collected into 13 x 100 mm borosilicate tubes with a Teflon-lined cap (catalog #60827-453, VWR, West Chester, PA). After addition of standards, lipids were extracted by the method of Bligh and Dyer. The extract was reduced to dryness using a Speed Vac. The dried residue was reconstituted in 0.2 ml of the starting mobile phase solvent for untargeted analysis, sonicated for 15 sec, then centrifuged for 5 minutes in a tabletop centrifuge before transfer of the clear supernatant to the autoinjector vial for analysis.

The lipids were separated by reverse phase LC using a Thermo Scientific Accucore Vanquish C18+ 2.1 (i.d.) x 150 mm column with 1.5 µm particles. The UHPLC used a binary solvent system at a flow rate of 0.26 mL/min with a column oven set to 55°C. Prior to injection of the sample, the column was equilibrated for 2 min with a solvent mixture of 99% Moble phase A1 (CH_3_CN/H_2_O, 50/50, v/v, with 5 mM ammonium formate and 0.1% formic acid) and 1% Mobile phase B1 (CH_3_CHOHCH_3_/CH_3_CN/H_2_O, 88/10/2, v/v/v, with 5 mM ammonium formate and 0.1% formic acid) and after sample injection (typically 10 μL), the A1/B1 ratio was maintained at 99/1 for 1.0 min, followed by a linear gradient to 35% B1 over 2.0 min, then a linear gradient to 60% B1 over 6 min, followed by a linear gradient to 100% B1 over 11 min., which held at 100% B1 for 5 min, followed by a 2.0 min gradient return to 99/1 A1/B1. The column was re-equilibrated with 99:1 A1/B1 for 2.0 min before the next run. Each sample was injected two times for analysis in both positive and negative modes. For initial full scan MS (range 300 to 2000 *m/z*) the resolution was set to 120,000 with a data-dependent MS^2^ triggered for any analyte reaching 3e6 or above signal. Data-dependent MS^2^ were collected at 30,000 resolution. Data was analyzed using Thermo Scientific’s Lipid Search 4.2 software.

### Statistical analysis

Statistical comparisons were made using a student t-test or analysis of variance (ANOVA) with post hoc Tukey or Bonferroni correction where appropriate. Statistical significance was defined as p<0.05. Data are presented as mean ± standard error of the mean. Fold change of ≥2 was considered significant. Pearson correlation coefficients were calculated using GraphPad Prism software.

## Results

### Hepatic steatosis exacerbates liver injury after warm partial hepatic ischemia reperfusion surgery

Starting at six weeks of age, male C57BL/6J mice were fed either a standard chow diet (non-steatotic) or a diet providing 42% of its calories as fat with 0.2% cholesterol diet for eight weeks (steatotic). Specific details regarding the two different diets are noted in Table 1 and Table 2. Mice on the steatotic diet gained significantly more weight and developed fatty liver after eight weeks on diet (Supplemental Figure 1). After eight weeks on diet, mice were subjected to either sham or IR surgery as detailed in the methods section (Figure 1A). After surgery, mice were recovered for 6 h, 24 h, or 72 h. As expected, plasma alanine transaminase activity (ALT), a marker of liver injury, was elevated in all mice undergoing IR surgery compared to sham at 6 and 24 h post-surgery. ALT was also significantly higher in mice with steatotic livers compared to non-steatotic livers at 6 h and 24 h post reperfusion (Figure 1B). Additionally, after IR surgery, liver expression of inflammatory markers including *Tnf* and *Il1b* was significantly increased compared to sham and significantly higher in steatotic liver compared to non-steatotic liver (Figure 1C, 1D). Following IR surgery, steatotic livers had more extensive areas of hepatic necrosis compared to non-steatotic liver (Figure 1E). These findings of increased liver injury and the temporal manifestation of this injury are consistent with our previous work in this model (Liss 2018 and 2020).

**Figure 1.**
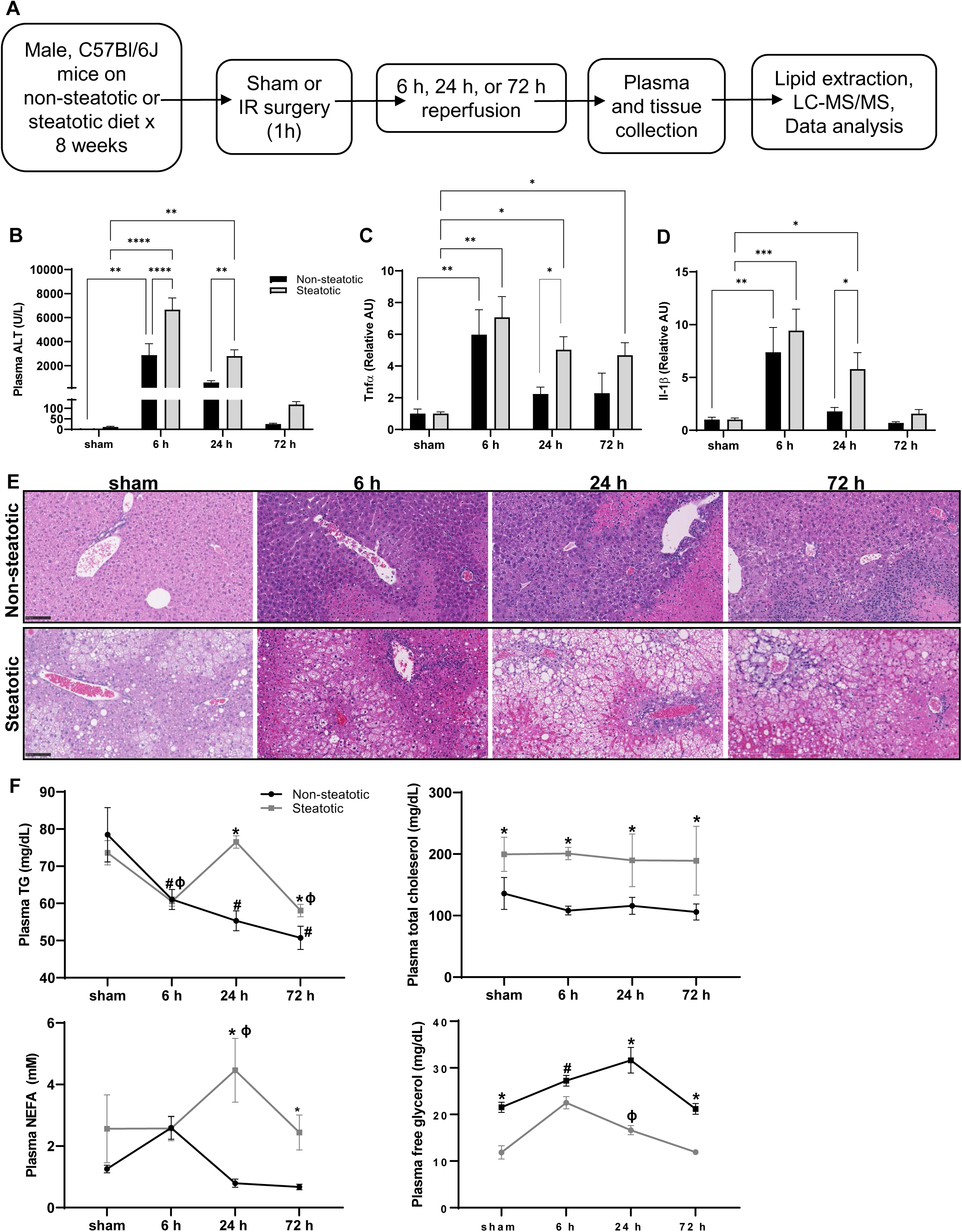
Mouse model of warm hepatic ischemia reperfusion injury, work flow, and indicators of liver injury. **A**. Schematic of experimental design. **B**. Plasma alanine aminotransferase (ALT) concentration following sham or IR surgery. **C** and **D**. Hepatic gene expression of inflammatory cytokines *Tnf* and *Il1b* following sham or IR surgery. **E**. Liver sections stained with hematoxylin and eosin following sham or IR surgery. **F**. Plasma lipid parameters following sham or IR surgery. Values are mean ± SEM. n = 4 per sham group, 8-10 per IR surgery group. Sham indicates sham surgery. 6 h, 24 h, 72 h indicates hours of reperfusion following IR surgery. *p<0.05, **p<0.01, p***<0.001, p****<0.0001. Black bars are chow fed mice and represent non-steatotic liver. Gray bars are 42% HF fed mice and represent steatotic liver. * indicates p<0.05 between non-steatotic and steatotic, # indicates p<0.05 between sham non-steatotic and non-steatotic IR at specified reperfusion time point. □ indicates p<0.05 between sham steatotic and steatotic IR at specified reperfusion time point.

**Table 1.** Dietary macronutrient composition

|  | % kcal |  |
| --- | --- | --- |
|  | 42% high fat diet,<br>Envigo TD.88137<br>(42% HF) | Standard chow diet,<br>PicoLab Rodent<br>Diet 20 5053<br>(Chow) |
| Protein | 15.2 | 24.6 |
| Carbohydrate | 42.7 | 62.1 |
| Fat | 42.0 | 13.2 |

**Table 2.**
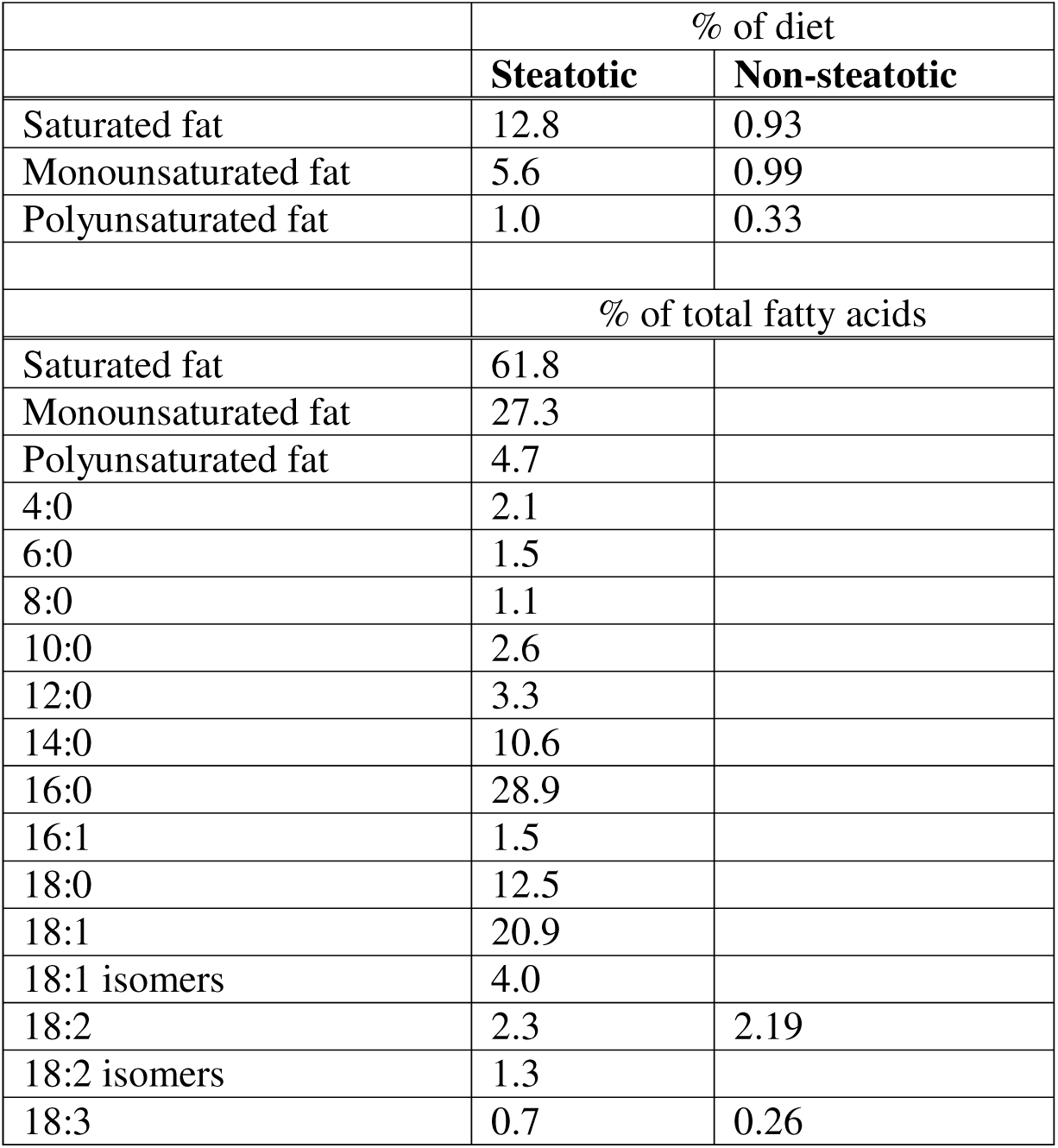
Dietary fatty acid profile

|  | % of diet |  |
| --- | --- | --- |
|  | **Steatotic** | **Non-steatotic** |
| Saturated fat | 12.8 | 0.93 |
| Monounsaturated fat | 5.6 | 0.99 |
| Polyunsaturated fat | 1.0 | 0.33 |
|  | % of total fatty acids |  |
| Saturated fat | 61.8 |  |
| Monounsaturated fat | 27.3 |  |
| Polyunsaturated fat | 4.7 |  |
| 4:0 | 2.1 |  |
| 6:0 | 1.5 |  |
| 8:0 | 1.1 |  |
| 10:0 | 2.6 |  |
| 12:0 | 3.3 |  |
| 14:0 | 10.6 |  |
| 16:0 | 28.9 |  |
| 16:1 | 1.5 |  |
| 18:0 | 12.5 |  |
| 18:1 | 20.9 |  |
| 18:1 isomers | 4.0 |  |
| 18:2 | 2.3 | 2.19 |
| 18:2 isomers | 1.3 |  |
| 18:3 | 0.7 | 0.26 |

To begin our evaluation of lipidomic changes following IRI, plasma lipid concentrations were quantified following sham and IR surgery (Figure 1F). We detected significant changes in plasma triglycerides (TG), free fatty acids (NEFA), and free glycerol between dietary groups and following IR surgery when compared to sham mice of each respective diet group. Specifically, plasma TG decreases following IR surgery at 6 h and 72 after reperfusion in both non-steatotic and steatotic diet fed mice. There were no changes to plasma total cholesterol concentrations following IR surgery in either diet groups. In steatotic mice, both NEFA and free glycerol increased in the plasma 24 h following IRI. In non-steatotic mice, there was a significant increase in plasma free glycerol following IR surgery at 6 h and a trend towards an increase in plasma NEFA at 6 h following surgery (p=0.14). Thus, in addition to liver inflammation and injury, there were dynamic changes in plasma lipid content in response to IR surgery in both non-steatotic and steatotic diet fed mice.

### Hepatic lipidomic profile following IRI

To fully characterize changes in hepatic lipid content following IR surgery, lipids were extracted from the median lobe (an ischemic lobe) from both non-steatotic and steatotic diet fed mice followed by LC-MS/MS. From the sham operated mice, a portion of the median lobe was also obtained for lipid extraction and analysis. Untargeted lipidomics identified 308 distinct lipid species in the following classes: triglyceride (TG), diglyceride (DG), ceramide (Cer), cardiolipin (CL), acylcarnitine (AcCa), phosphatidylcholine (PC), phosphatidic acid (PA), phosphatidylglycerol (PG), phosphatidylinositol (PI), phosphatidylserine (PS), phosphatidylethanolamine (PE), lysophosphatidylcholine (LPC), lysophosphatidylethanolamine (LPE), lysophosphatidylserine (LPS), lysophosphatidylinositol (LPI), coenzyme Q (CoQ), hexosylceramide (HexCer), and endocannabinoids. A complete list of lipid species is delineated in Table 3.

**Table 3.** Lipid classes identified in liver following ischemia reperfusion injury

| Lipid class | Lipid species |
| --- | --- |
| Glycerolipids | DG(16:1_18:2), DG(16:1_18:1), DG(18:3_18:2), DG(18:2_18:2), DG(18:1_18:2), DG(18:1_18:1), DG(18:2_20:4), DG(16:0_22:6), DG(18:1_20:4), DG(18:0_20:4), DG(20:1_18:2), DG(20:1_18:1), DG(18:2_22:6), DG(22:5_18:2), DG(18:1_22:6), DG(18:1_22:5), DG(18:0_22:6), DG(22:1_18:2)<br>TG(12:0_14:0_14:0), TG(16:0_12:0_14:0), TG(16:0_14:0_14:0), TG(16:0_12:0_18:2), TG(18:0_10:4_20:5), TG(16:1_14:0_18:2), TG(16:0_16:1_16:1), TG(16:0_14:0_18:1), TG(16:0_16:0_16:0), TG(15:0_16:1_18:2), TG(15:0_16:0_18:2), TG(15:0_16:0_18:1), TG(14:0_18:2_18:3), TG(16:0_16:1_18:3) TG(16:1_16:1_18:2), TG(16:0_16:1_18:2) TG(16:0_16:1_18:1), TG(16:0_16:0_18:1), TG(15:0_18:2_18:3), TG(16:1_17:1_18:2) TG(16:1_17:1_18:1), TG(16:0_17:1_18:1) TG(16:0_17:0_18:1), TG(18:4_16:1_18:2) TG(16:1_18:2_18:3), TG(16:1_18:2_18:2) TG(16:0_18:2_18:3), TG(16:0_18:2_18:2) TG(16:0_18:1_18:3), TG(16:0_18:1_18:2) TG(16:0_18:1_18:1), TG(18:3_17:1_18:2) TG(18:2_17:1_18:2), TG(18:1_17:1_18:2) TG(17:0_18:1_18:2), TG(18:3_18:3_18:3) TG(17:0_18:1_18:1), TG(18:3_18:2_18:3) TG(16:0_18:1_19:0), TG(18:3_18:2_18:2) TG(16:0_18:2_20:5), TG(18:1_18:2_18:3) TG(18:2_18:2_18:2), TG(18:1_18:2_18:2) TG(18:1_18:1_18:3), TG(18:1_18:1_18:2) TG(18:1_18:1_18:1), TG(18:0_18:1_18:1) TG(15:0_18:2_22:6), TG(20:0_16:0_18:1) TG(15:0_18:1_22:6), TG(18:1_17:1_20:4) TG(19:1_18:1_18:2), TG(18:3_18:2_20:5) TG(19:0_18:1_18:2), TG(20:5_18:2_18:2) TG(19:0_18:1_18:1), TG(16:0_18:2_22:6) TG(16:0_18:1_21:0), TG(16:0_18:2_22:5) TG(16:0_18:1_22:6), TG(18:1_18:2_20:4) TG(16:0_18:1_22:5), TG(18:1_18:1_20:4) TG(18:1_18:2_20:2), TG(20:1_18:1_18:2) TG(18:1_18:1_20:2), TG(20:1_18:1_18:1) TG(18:2_17:1_22:6), TG(16:0_18:1_22:1) TG(22:5_17:1_18:2), TG(17:0_18:2_22:6) TG(16:0_18:1_22:0), TG(17:0_18:1_22:6) TG(18:3_18:3_22:6), TG(18:4_18:2_22:6) TG(18:3_18:2_22:6), TG(20:5_18:2_20:4) TG(16:0_20:5_22:6), TG(18:2_18:2_22:6) TG(18:1_18:3_22:6), TG(18:1_18:2_21:0) TG(22:5_18:2_18:2), TG(18:1_18:2_22:6) TG(18:1_18:1_21:0), TG(18:1_18:2_22:5) TG(18:1_18:1_22:6), TG(18:0_18:1_22:6) TG(18:1_18:2_22:4), TG(18:1_18:2_22:3) TG(22:1_18:2_18:2), TG(20:0_18:1_20:4) TG(18:1_18:2_22:1), TG(18:1_18:1_22:1) TG(18:1_18:1_22:0), TG(16:0_18:1_24:0) TG(18:3_20:5_22:6), TG(20:5_18:2_22:6) TG(18:2_20:4_22:6), TG(16:0_22:6_22:6) TG(16:0_22:5_22:6), TG(22:5_18:2_20:4) TG(18:0_20:4_22:6), TG(16:0_22:5_22:5) TG(18:1_18:2_23:0), TG(20:1_18:2_22:6) TG(20:1_18:2_22:5), TG(20:1_18:1_22:6) TG(20:0_18:1_22:6), TG(24:1_18:2_18:2) TG(18:1_18:2_24:1), TG(18:1_18:1_24:1) TG(18:1_18:1_24:0), TG(18:2_22:6_22:6) TG(22:5_18:2_22:6), TG(18:1_22:6_22:6) TG(18:1_22:5_22:6), TG(18:1_22:5_22:5) TG(22:1_18:2_22:6), TG(26:1_18:1_18:1) TG(20:2_22:6_22:6) |
| Glycerophospholipids | PA(18:0_20:4)<br>PC(16:1_16:1), PC(16:0_16:1), PC(16:0_16:0), PC(15:0_18:2), PC(16:1_18:3), PC(14:0_20:4), PC(16:1_18:1), PC(16:0_18:2), PC(16:0_18:1) , PC(18:0_16:0), PC(15:0_20:4), PC(17:1_18:2), PC(17:0_18:2), PC(17:0_18:1), PC(16:1_20:5), PC(18:3_18:2), PC(16:0_20:5) , PC(18:2_18:2), PC(16:0_20:4), PC(18:1_18:2), PC(18:1_18:1) , PC(18:0_18:2), PC(18:0_18:1), PC(17:0_20:4),, PC(19:0_18:2), PC(16:1_22:6), PC(18:2_20:4), PC(16:0_22:6), |
|  | PC(18:1_20:4), PC(18:1_20:3), PC(18:0_20:4), PC(18:0_20:3),<br>PC(20:0_18:2) , PC(19:0_20:4), PC(18:0_22:6), PC(18:0_22:4),<br>PC(20:0_20:4), PC(20:0_20:3), PC(20:4_22:6), PC(20:1_22:6),<br>PC(20:0_22:6)<br>PE(16:0_16:0), PE(16:1_18:2), PE(16:0_18:1), PE(18:0_16:0) ,<br>PE(16:0p_20:4), PE(18:0p_18:2), PE(17:0_18:2), PE(18:3_18:2),<br>PE(16:1_20:4), PE(16:0_20:5), PE(18:2_18:2), PE(16:0_20:4),<br>PE(18:1_18:2), PE(18:1_18:1), PE(18:0_18:2), PE(18:0_18:1),<br>PE(16:0p_22:6), PE(15:0_22:6), PE(16:0p_22:5), PE(18:0p_20:4),<br>PE(17:0_20:4) , PE(16:1_22:6), PE(18:2_20:4), PE(16:0_22:6),<br>PE(18:1_20:4), PE(18:0_20:4), PE(20:0_18:2), PE(18:1e_22:6),<br>PE(18:0p_22:6), PE(18:0p_22:5), PE(19:0_20:4), PE(18:3_22:6),<br>PE(18:2_22:6), PE(18:1_22:6), PE(18:0_22:6), PE(20:0_22:6),<br>PG(22:6_22:6)<br>PI(16:1_18:2), PI(16:0_18:2), PI(16:0_20:5), PI(18:2_18:2),<br>PI(16:0_20:4), PI(18:1_18:2), PI(16:0_20:3), PI(18:0_18:2),<br>PI(18:1_18:1), PI(18:0_18:1), PI(17:0_20:4), PI(18:2_20:4),<br>PI(16:0_22:6), PI(18:0_20:5), PI(18:1_20:4), PI(18:0_20:4),<br>PI(18:0_20:3), PI(19:0_20:4), PI(20:4_20:4), PI(18:0_22:6),<br>PI(18:0_22:5), PI(18:0_22:4), PI(20:0_20:4)<br>PS(16:0_18:2), PS(16:0_20:5), PS(16:0_20:4), PS(18:0_18:2),<br>PS(18:0_18:1), PS(16:0_22:6), PS(18:1_20:4), PS(18:0_20:5),<br>PS(18:0_20:4), PS(20:4_20:4), PS(18:1_22:6), PS(18:0_22:6),<br>PS(18:0_22:4), PS(20:4_22:6)<br>LPC(16:1), LPC(16:0), LPC(17:0), LPC(18:3), LPC(18:2), LPC(18:1),<br>LPC(18:0), LPC(19:0), LPC(20:5), LPC(20:4), LPC(20:3), LPC(20:2),<br>LPC(20:1), LPC(22:6)<br>LPE(16:1), LPE(16:0), LPE(18:2), LPE(18:1), LPE(18:0), LPE(22:6)<br>LPI(16:0), LPI(18:2), LPI(18:1), LPI(18:0), LPI(20:4), LPI(20:3) LPS(18:0)<br>CL(18:1_16:0_16:0_18:2), CL(18:2_16:1_16:1_20:3),<br>CL(18:2_16:1_18:2_18:1), CL(18:2_18:1_16:1_18:1),<br>CL(18:3_18:2_18:2_18:2), CL(18:2_18:2_18:2_18:2),<br>CL(18:2_18:2_18:2_18:1), CL(18:2_18:2_18:2_22:6) |
| Sphingolipids | Cer(d18:1_22:0), Cer(d18:1_23:0), Cer(d18:1_24:1), Cer(d18:1_24:0),<br>Hex1Cer(d18:1_22:0), Hex1Cer(d18:1_23:0), Hex1Cer(d18:1_24:0) |
| Other | AEA(16:0), AcCa(18:2), AcCa(18:0), AcCa(20:4), AcCa(20:0), AcCa(22:1),<br>AcCa(22:0), AcCa(24:1), Co(Q8), Co(Q9), Co(Q10) |

The principal component analysis (PCA) plot highlighted differences and similarities between non-steatotic and steatotic diet fed mice and indicated that the greatest source of variation was between the two diet groups (Figure 2A). Indeed, the non-steatotic and steatotic livers formed two distinct clusters. Within the non-steatotic group, the sham and 6 h reperfusion were very similar while 24 h and 72 h reperfusion time points clustered together. In contrast, within the steatotic group, the sham, 6 h, and 24 h were similarly clustered, while the 72 h reperfusion time point formed a distinct subgroup with wide variability within its subgroup.

**Figure 2.**
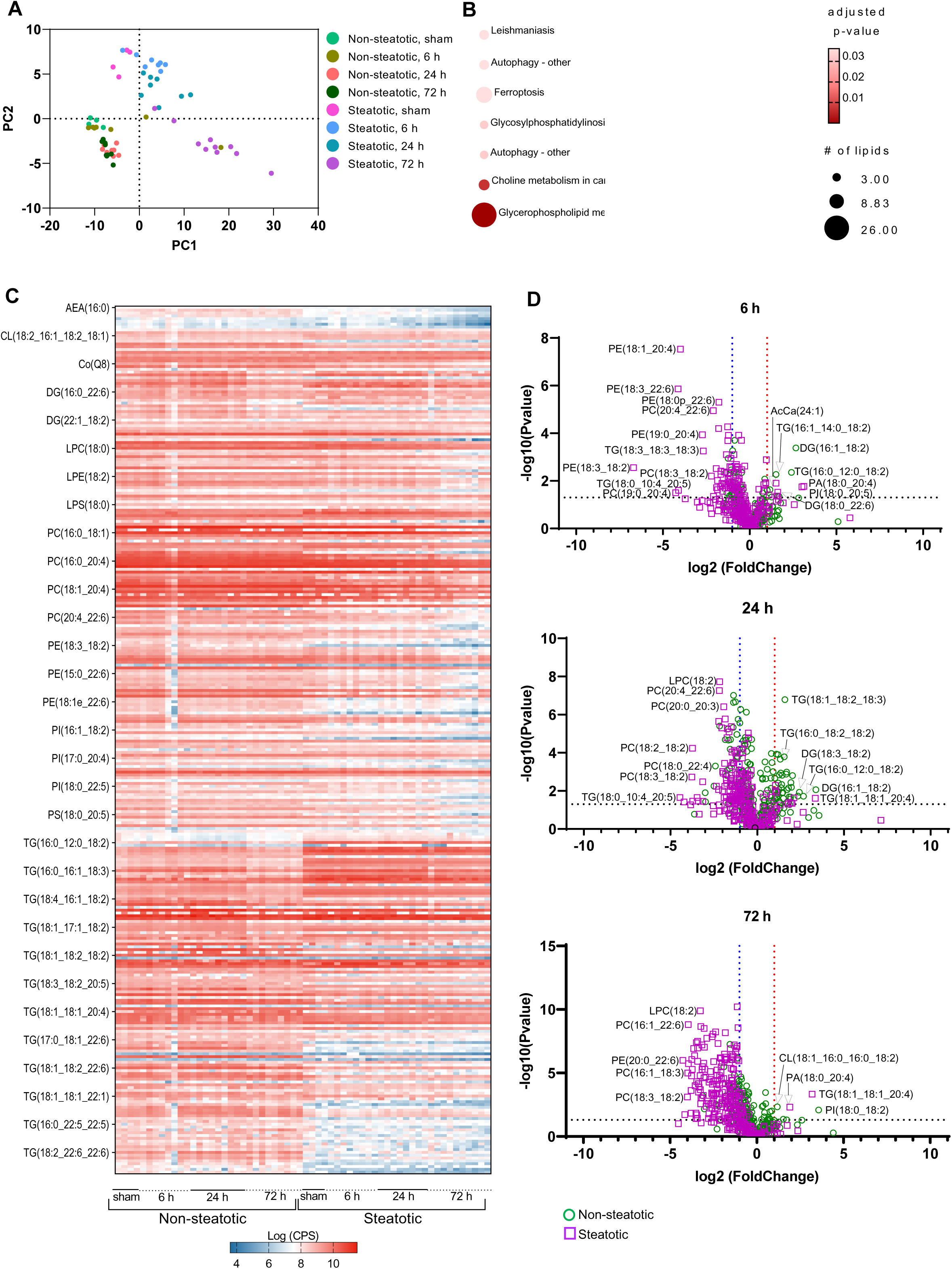
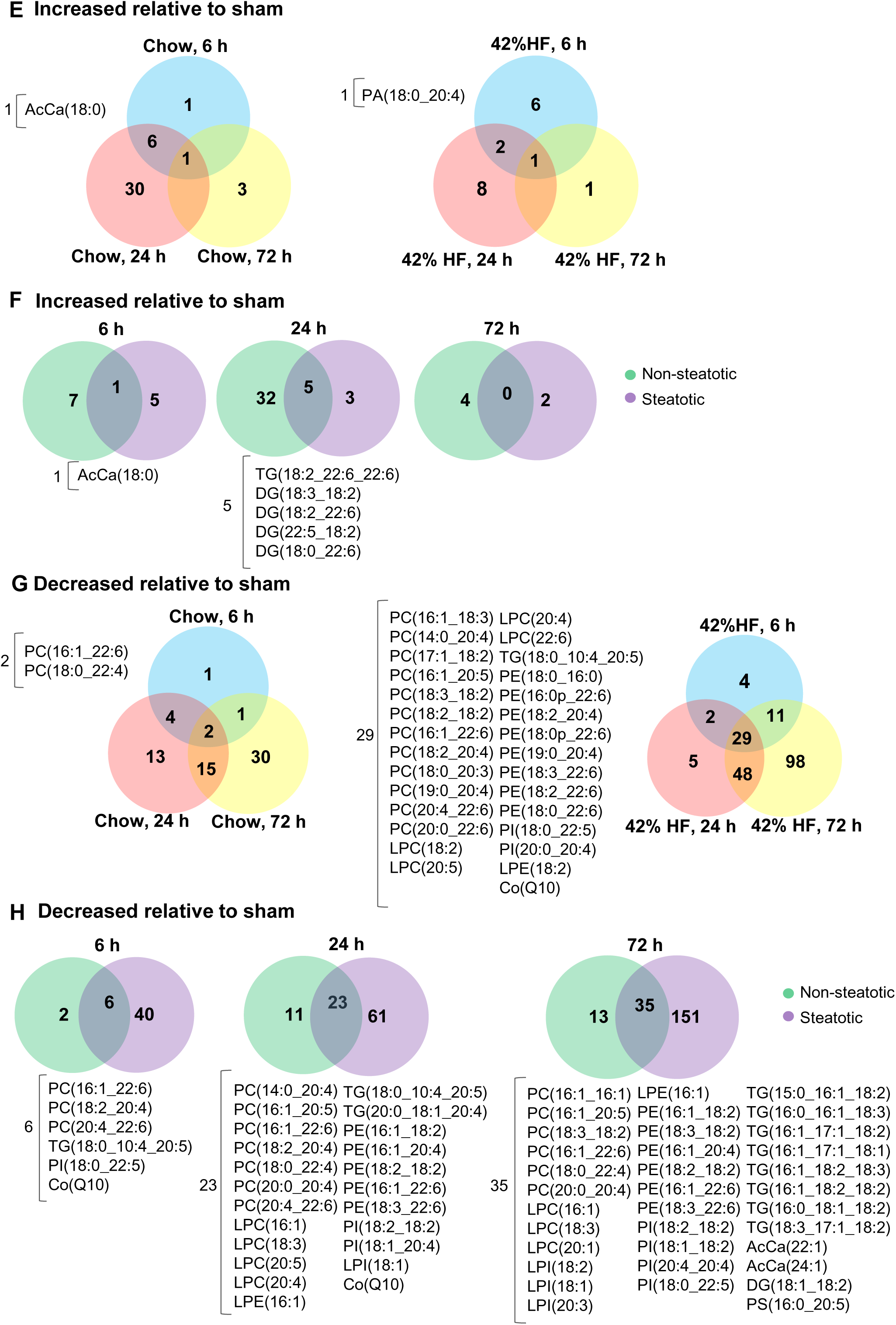
Lipidomic profile analysis. **A**. Principal component analysis plot of the lipidomic profiles of sham operated and IR surgery groups in non-steatotic and steatotic diet fed groups. **B**. LIPEA pathway analysis indicating enriched pathways in the KEGG database. **C**. Heat map indicating LogCPS of all identified lipid species. **D**. Volcano plots at specified reperfusion time points following IR surgery. The Log2 fold change is on the x axis, and the P value is converted to the – log10 scale is on the y axis. Fold change is relative to respective diet sham. Dashed line indicates threshold for significant p-value (-Log10(pvalue)>1.3). Non-steatotic, green circles. Steatotic, purple square. **E, F, G, H**. Venn diagram showing the overlap and differences of differentially expressed lipids in the respective groups. The numbers shown in overlapping areas illustrate the number of lipids commonly differentially expressed. n = 4 per sham group, 8-10 per IR surgery group. Sham indicates sham surgery. 6 h, 24 h, 72 h indicates hours of reperfusion following IR surgery.

Pathway enrichment analysis performed using lipid pathway enrichment analysis (LIPEA) indicated that Glycerophospholipid metabolism was most dramatically altered and was significantly associated with the set of lipids that were identified following IRI (Figure 2B). After Bonferonni correction, Choline metabolism in cancer was also identified as significantly changed with IRI (adjusted p-value 0.014). Log transformed counts per second (CPS) values of all identified lipids highlighted the changes of lipid metabolites following IR surgery and demonstrated a clear distinction between steatotic and non-steatotic liver in both sham operated animals and following IR surgery. The most notable differences were in TG species and glycerophospholipids (Figure 2C).

We then looked at individual lipid species and their fold change over their respective diet shams at all reperfusion time points (Figure 2D). Compared to non-steatotic livers (green circles), steatotic livers (purple squares) contained more lipids that were decreased relative to sham at all reperfusion time points. While most lipid species in non-steatotic livers either remained elevated or returned to sham levels at 72 h, many lipid species were significantly decreased relative to sham in steatotic livers even 72 h after reperfusion. A complete list of significantly increased or decreased lipid species at each reperfusion time point is delineated in Supplemental Table 1.

We then compared lipids with altered abundances within and between diet groups (Figure 2E, F, G, H). Relative to sham, only a few of the identified lipids increased in both non-steatotic and steatotic liver following IRI (Figure 2E). In non-steatotic liver, AcCa(18:0) was the only lipid that increased at all reperfusion time points. In steatotic liver, PA(18:0_20:4) was increased at all time points (Figure 2E). Comparison of non-steatotic and steatotic liver at individual reperfusion time points identified only six shared lipid species that increased relative to sham (Figure 2F).

We next looked at lipids that decreased relative to sham following IRI. Strikingly, in both steatotic and non-steatotic liver, the number of lipids that were significantly decreased relative to sham increased with reperfusion time, and the number of lipid species conforming to this pattern, was more pronounced in steatotic liver (Figure 2G). In non-steatotic liver, there were only two lipids that were decreased at all reperfusion time points. In steatotic liver, there were 29 lipid species that were decreased at all reperfusion time points (Figure 2G). At all reperfusion time points, there were more lipids that decreased relative to sham in steatotic liver compared to non-steatotic liver (Figure 2H). Collectively, these data highlight the distinct differences between non-steatotic and steatotic liver and demonstrate dynamic changes in lipid composition following IRI, which was most notable for a dramatic decrease in multiple lipid classes.

### Comparison of TG and DG species

Triglyceride (TG) comprises the primary storage lipid and is synthesized by sequential acylation of fatty acids to a glycerol backbone. As expected, mice fed a steatotic diet had higher total TG in the liver compared to non-steatotic diet fed mice at baseline (sham groups; Figure 3A). This was consistent with the presence of steatosis on histological exam of liver from mice fed a steatotic diet (Supplemental Figure 1). In non-steatotic liver, there was an increase in total TG following IRI (relative to sham) at 24 h (Figure 3A) and multiple TG species were significantly increased relative to sham at 24 h (Figure 3B, red circles). In steatotic liver, there was a decrease in total TG following IRI (relative to sham) at 24 h and 72 h (Figure 3A) and multiple individual TG species were decreased relative to sham at 24 h (Figure 3C, blue circles) and 72 h. Notably, at 24 h reperfusion, non-steatotic diet fed mice had a significantly higher total TG content in the liver than mice fed the steatotic diet (Figure 3A).

**Figure 3.**
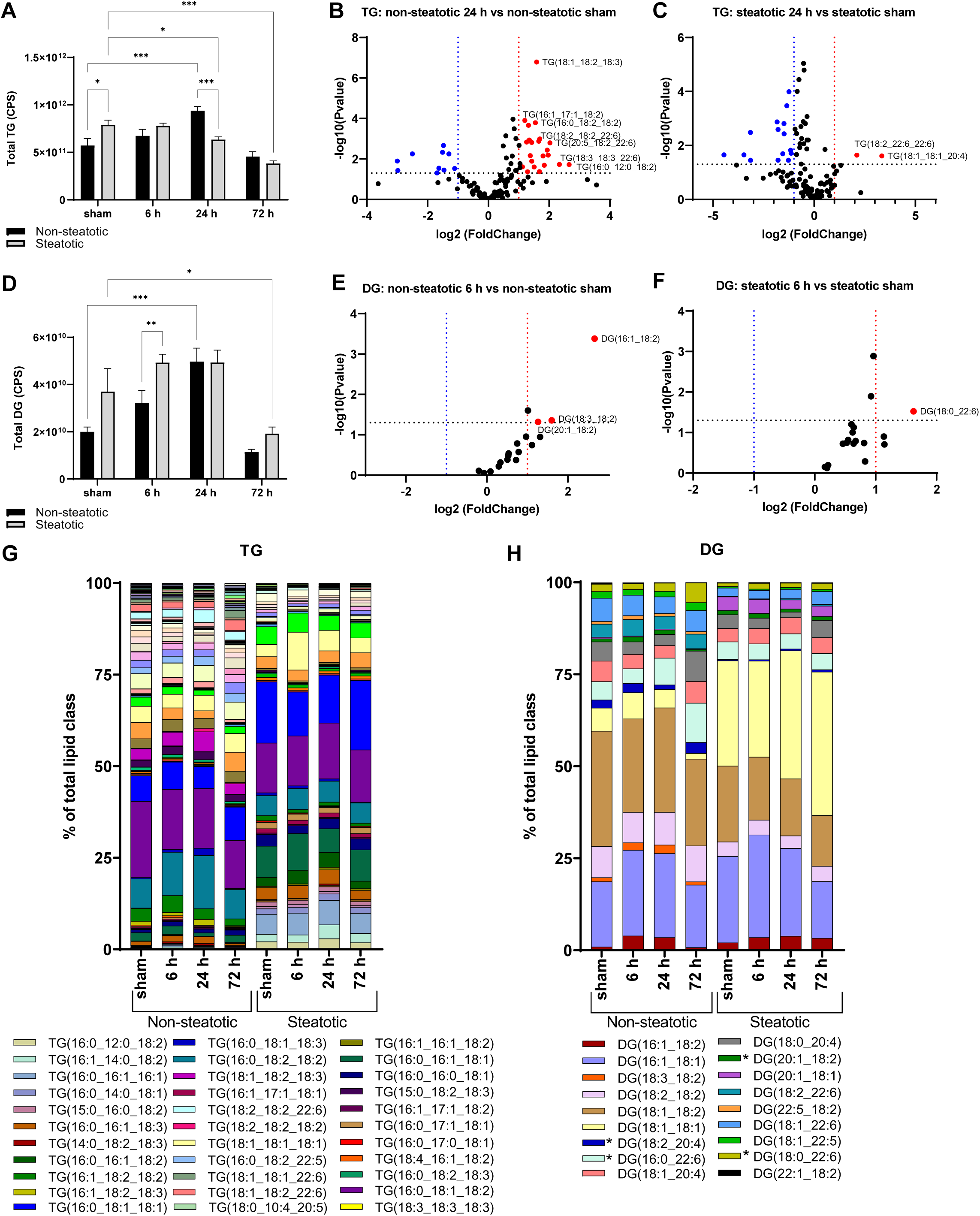
Analysis of glycerolipids. **A**. Quantification of total triglyceride (TG) following sham or IR surgery. **B**. Volcano plot of specific TG species comparing 24 h reperfusion relative to sham in non-steatotic liver. **C**. Volcano plot of specific TG species comparing 24 h reperfusion relative to sham in steatotic liver. **D**. Quantification of total diglyceride (DG) following sham or IR surgery. **E**. Volcano plot of specific DG species comparing 6 h reperfusion and non-steatotic sham. **F**. Volcano plot of specific DG species comparing 6 h reperfusion and steatotic sham. **G**. Specific TG species represented as a percent of total TG lipids following sham or IR surgery. A complete legend is provided in Supplemental Figure 1. **H**. Specific DG species represented as a percent of total DG lipids following sham or IR surgery. * in legend indicates significant difference from respective diet sham. Values are mean ± SEM. n = 4 per sham group, 8-10 per IR surgery group. Sham indicates sham surgery. 6 h, 24 h, 72 h indicates hours of reperfusion following IR surgery. *p<0.05, **p<0.01, p***<0.001, p****<0.0001. For volcano plots, lipids that are significantly increased relative to sham are represented in red, and those significantly decreased relative to sham are represented in blue. Black bars are chow fed mice and represent non-steatotic liver. Gray bars are 42% HF fed mice and represent steatotic liver.

Following a pattern similar to total TG, total diglyceride (DG) increased significantly in non-steatotic liver at 24 h reperfusion compared to sham and returned to baseline at 72 h (Figure 3D). In contrast, total DG in the steatotic liver decreased significantly at 72 h compared to steatotic sham. There was a trend towards a higher total DG content in steatotic liver (compared to non-steatotic liver) in sham operated animals and at 72 h reperfusion (p = 0.06 and 0.18, respectively). Total DG was significantly higher in steatotic liver compared to non-steatotic liver at 6 h. Although steatotic liver had higher total DG content, only one specific DG species was significantly increased from sham at 6 h. In contrast, in non-steatotic liver, three different DG species were significantly increased relative to chow sham at 6 h (Figure 3E, F).

We next looked at each TG and DG species as a percent of total TG and DG, respectively (Figure 3G, H), which highlighted the compositional changes within a lipid class with reperfusion. In non-steatotic liver, as a percent of total TGs and DGs, seven TGs and four DGs were significantly changed relative to non-steatotic sham (Figure 3G, H, Supplemental Figure 3). In steatotic liver, as a percent of total TGs and DGs, only TG(18:1_18:1_20:4) and none of the detected DG species was significantly changed relative to steatotic sham (Figure 3G, H, Supplemental Figure 3). Lipid species in legends marked with an asterisk are significantly different compared to corresponding diet shams.

In summary, there was an increase in multiple TG and DG species following IRI in non-steatotic liver which then returned to baseline levels at 72 h after reperfusion. However, in steatotic liver, total TG or DG content did not increase compared to sham controls. Instead, in steatotic liver, total DG and TG content decreased at 72 h relative to sham. Furthermore, when compared to steatotic liver, non-steatotic liver demonstrated more compositional fluctuations with IRI while the percentage of each TG and DG species remained relatively similar with IRI in steatotic liver.

### Comparison of phospholipids

Phospholipids are important components of cellular membranes that impact membrane biophysical properties. In non-steatotic liver, total phosphatidylcholine (PC) was significantly decreased following IRI at all time points compared to non-steatotic sham (Figure 4A). In non-steatotic liver, total phosphatidylinositol (PI) decreased at 6 h compared to sham, but returned to baseline at 24 h and 72 h (Figure 4G). There were no significant changes in total phosphatidylethanolamine (PE) or phosphatidylserine (PS) with IRI in non-steatotic liver (Figure 4D, J). In the steatotic liver, total PC, PE, PI and PS were significantly decreased following IRI compared to steatotic sham. This was most dramatic at the 72 h reperfusion time point (Figure 4A, D, G, J).

**Figure 4.**
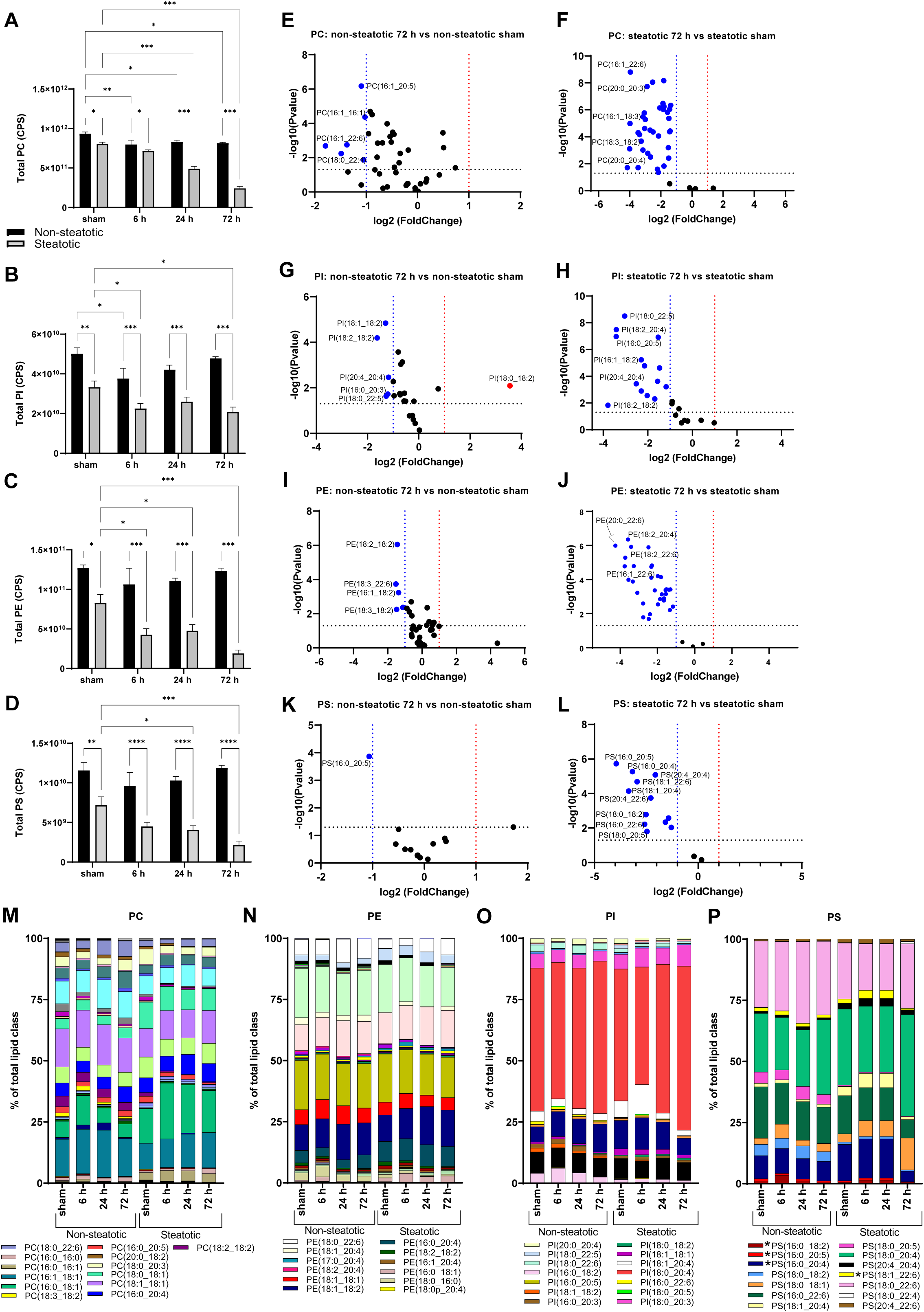
Analysis of glycerophospholipids. **A**. Quantification of total phosphatidylcholine (PC) following sham or IR surgery. **B**. Volcano plot of specific PC species comparing 72 h reperfusion and non-steatotic sham. **C**. Volcano plot of specific PC species comparing 72 h reperfusion and steatotic sham. **D**. Quantification of total phosphatidylethanolamine (PE) following sham or IR surgery. **E**. Volcano plot of specific PE species comparing 72 h reperfusion and non-steatotic sham. **F**. Volcano plot of specific PE species comparing steatotic 72 h reperfusion and sham. **G**. Quantification of total phosphatidylinositol (PI) following sham or IR surgery. **H**. Volcano plot of specific PI species comparing non-steatotic 72 h reperfusion and sham. **I**. Volcano plot of specific PI species comparing steatotic 72 h reperfusion and sham. **J**. Quantification of total phosphatidylserines (PS) following sham or IR surgery. **K**. Volcano plot of specific PS species comparing non-steatotic 72 h reperfusion and sham. **L**. Volcano plot of specific PS species comparing steatotic 72 h reperfusion and sham. Specific PC [**M**], PE [**N**], PI [**O**] species represented as a percent of total PC lipids following sham or IR surgery. A complete legend is provided in Supplemental Figure 1. *in legend indicates significant difference from respective diet sham. **P**. Specific PS species represented as a percent of total PS lipids following sham or IR surgery. Values are mean ± SEM. n = 4 per sham group, 8-10 per IR surgery group. Sham indicates sham surgery. 6 h, 24 h, 72 h indicates hours of reperfusion following IR surgery. *p<0.05, **p<0.01, p***<0.001, p****<0.0001. For volcano plots, lipids that are significantly increased relative to sham are represented in red, and those significantly decreased relative to sham are represented in blue. Black bars are chow fed mice and represent non-steatotic liver. Gray bars are 42% HF fed mice and represent steatotic liver.

There were significant differences between non-steatotic and steatotic liver in sham animals as well as all reperfusion time points for PC, PE, PI, and PS (Figure 4A, D, G, J). The greatest difference between non-steatotic and steatotic liver was at the 72 h time point. To evaluate the 72 h reperfusion time point more closely, we looked at fold change of specific lipid species relative to the respective diet sham (Figure 4). Strikingly, only six PC species were significantly decreased relative to sham in non-steatotic liver while the majority of PC species were significantly decreased relative to sham in steatotic liver (Figure 4B, C). Both PE and PI followed a similar pattern (Figure 4E, F, H, I). Of note, one PI species (PI(18:0_18:2)) was significantly increased relative to sham in non-steatotic liver, while this same lipid was significantly decreased in steatotic liver at 72 h. One PS species was significantly changed relative to chow sham at 72 h reperfusion, while the majority of PS species were significantly decreased relative to sham in steatotic liver at 72 h reperfusion (Figure 4K, L).

We next looked at each individual phospholipid species as a percent of the total lipid class (Figure 4M, N, O, P). In non-steatotic liver, the proportion of twenty PC species significantly changed relative to the chow sham in at least one reperfusion time point. In steatotic liver, the proportion of eight PC species changed relative to steatotic sham in at least one reperfusion time point (Figure 4M). In non-steatotic liver, the proportion of fourteen PE species changed relative to chow sham in at least one reperfusion time point. In steatotic liver, the proportion of twelve PE species changed relative to chow sham in at least one reperfusion time point (Figure 4N). In non-steatotic liver, the proportion of five PI species changed relative to non-steatotic sham in at least one time point. In steatotic liver, the proportion of ten PI species changed in at least one time point compared to sham (Figure 4O). The proportion of only one PS species was significantly different than sham in non-steatotic liver. The proportion of four PS species in steatotic liver were significantly altered from sham in at least one time point (Figure 4P).

Together, these data indicate that there was a global disruption in phospholipid content marked by a dramatic decrease in most phospholipid species in steatotic liver following IRI. Additionally, while the total phospholipid content did not markedly fluctuate in non-steatotic liver following IRI, we noted significant shifts in lipid class composition.

### Comparison of lysoglycerophospholipids

We next examined changes in LPC, LPE, and LPI following IRI in non-steatotic and steatotic liver (Figure 5). The lysoglycerophospholipids followed a pattern similar to the phospholipids in response to IRI. In non-steatotic liver, total LPC and LPI decreased with IRI compared to non-steatotic sham (Figure 4A, G), but there was no significant change in total LPE with IRI (Figure 4D). In steatotic liver, total LPC and LPE decreased with IRI compared to steatotic sham, but there was no significant change in total LPI with IRI (Figure 4A, D, G).

**Figure 5.**
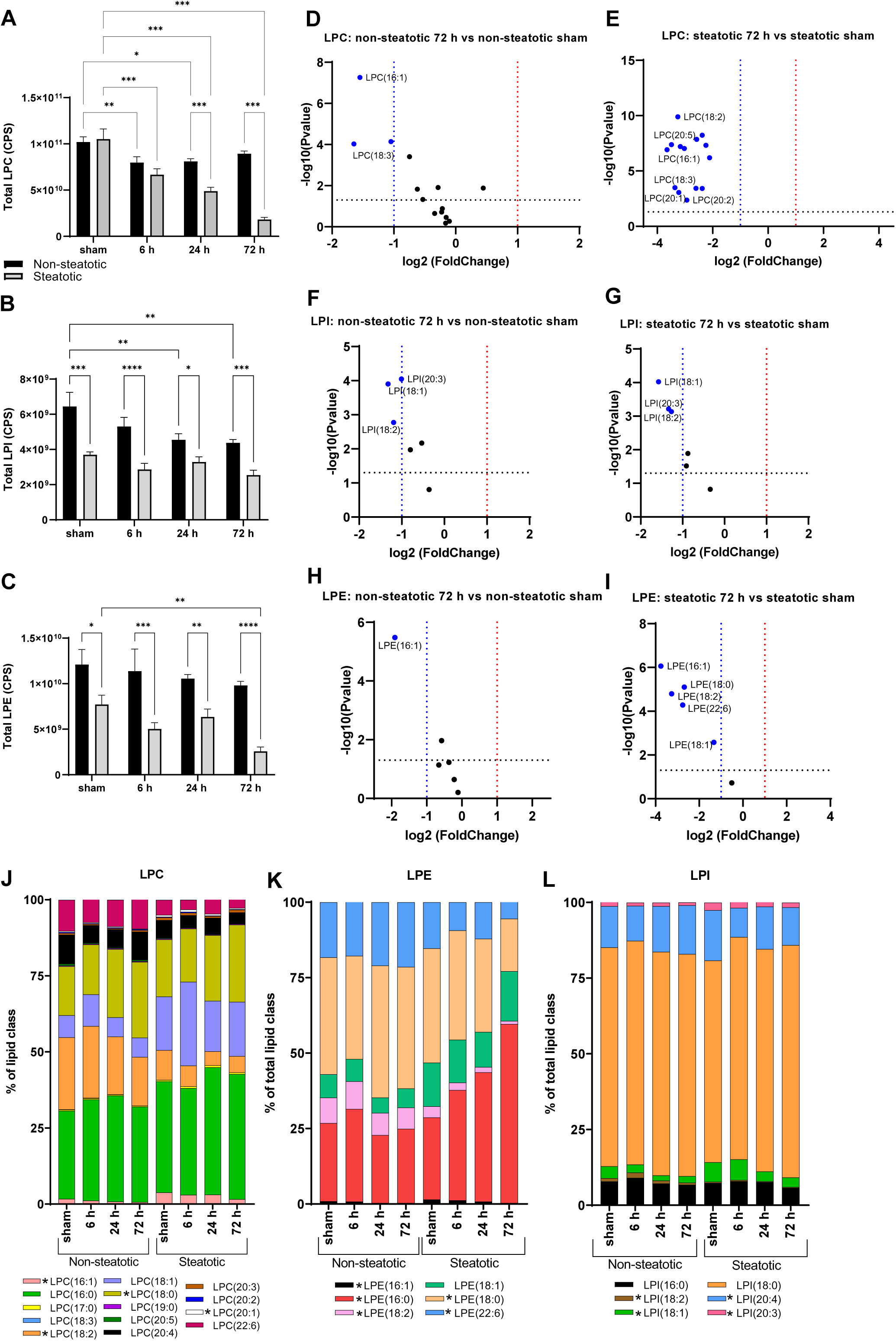
Analysis of lysophospholipids. **A**. Quantification of total lysophosphatidylcholine (LPC) following sham or IR surgery. **B**. Volcano plot of specific LPC species comparing non-steatotic 72 h reperfusion and sham. **C**. Volcano plot of specific LPC species comparing steatotic 72 h reperfusion and sham. **D**. Quantification of total lysophosphatidylethanolamine (LPE) following sham or IR surgery. **E**. Volcano plot of specific LPE species comparing non-steatotic 72 h reperfusion to sham. **F**. Volcano plot of specific LPE species comparing steatotic 72 h reperfusion and sham. **G**. Quantification of total lysophosphatidylinositol (LPI) following sham or IR surgery. **H**. Volcano plot of specific LPI species comparing non-steatotic 72 h reperfusion to sham. **I**. Volcano plot of specific LPI species comparing steatotic 72 h reperfusion and sham. **J**. Specific LPC species represented as a percent of total LPC lipids following sham or IR surgery. *in legend indicates significant difference from respective diet sham. **K**. Specific LPE species represented as a percent of total LPE lipids following sham or IR surgery. L. Specific LPI species represented as a percent of total LPI lipids following sham or IR surgery. Values are mean ± SEM. n = 4 per sham group, 8-10 per IR surgery group. Sham indicates sham surgery. 6 h, 24 h, 72 h indicates hours of reperfusion following IR surgery. *p<0.05, **p<0.01, p***<0.001, p****<0.0001. For volcano plots, lipids that are significantly increased relative to sham are represented in red, and those significantly decreased relative to sham are represented in blue. Black bars are chow fed mice and represent non-steatotic liver. Gray bars are 42% HF fed mice and represent steatotic liver.

Comparing non-steatotic and steatotic liver, total LPE and LPI were significantly different in sham and all reperfusion time points, with higher levels in non-steatotic liver compared to steatotic liver (Figure 5D, G). For LPC, there was a significant difference between non-steatotic and steatotic liver in sham at 24 and 72 h (Figure 5A). We again examined specific lipid species more closely at the 72 h reperfusion time point. For both LPC and LPE, there were strikingly more individual lipid species decreased relative to sham in steatotic liver compared to non-steatotic liver (Figure 5B, C, E, F). For LPI, in both non-steatotic and steatotic livers, three specific LPI species were significantly decreased relative to their respective diet sham group (Figure 5H, I).

To evaluate the relative compositional shift of specific lipid classes with IR injury, we examined lipid species as a percentage of the total for that particular class (Figure 5J, K, L). Of particular note, in non-steatotic liver, the proportion of LPE(16:0) did not change significantly with IRI at any reperfusion time point, but there was a significant and progressive increase in the proportion of LPE(16:0) following IR in steatotic liver.

In all, these data indicate that similar to phospholipids, essentially all detected LPC and LPE species were significantly decreased in the steatotic liver following IRI. While some of the LPC and LPE species were decreased also in non-steatotic liver, this was less pronounced compared to steatotic liver. Interestingly, the proportion of each lipid species as a percent of the total class did not have many prominent changes with the exception of LPE(16:0), which exhibited a dramatic increase in steatotic liver following IR, but no change in non-steatotic liver.

### Comparison of mitochondrial lipids

We next examined changes in lipid species closely associated with mitochondrial function or enriched in mitochondrial membranes (Figure 6). Acylcarnitines are synthesized to facilitate transport of fatty acyl groups across the inner mitochondrial membrane to the matrix for β-oxidation. In non-steatotic liver, total acylcarnitines (AcCa) increased with IRI compared to non-steatotic sham and AcCa(18:0) was increased at all reperfusion time points (Figure 6A, B). In steatotic liver, total AcCa did not change following IRI relative to steatotic sham, but multiple individual AcCa species were significantly decreased following IRI (Figure 6A, C).

**Figure 6.**
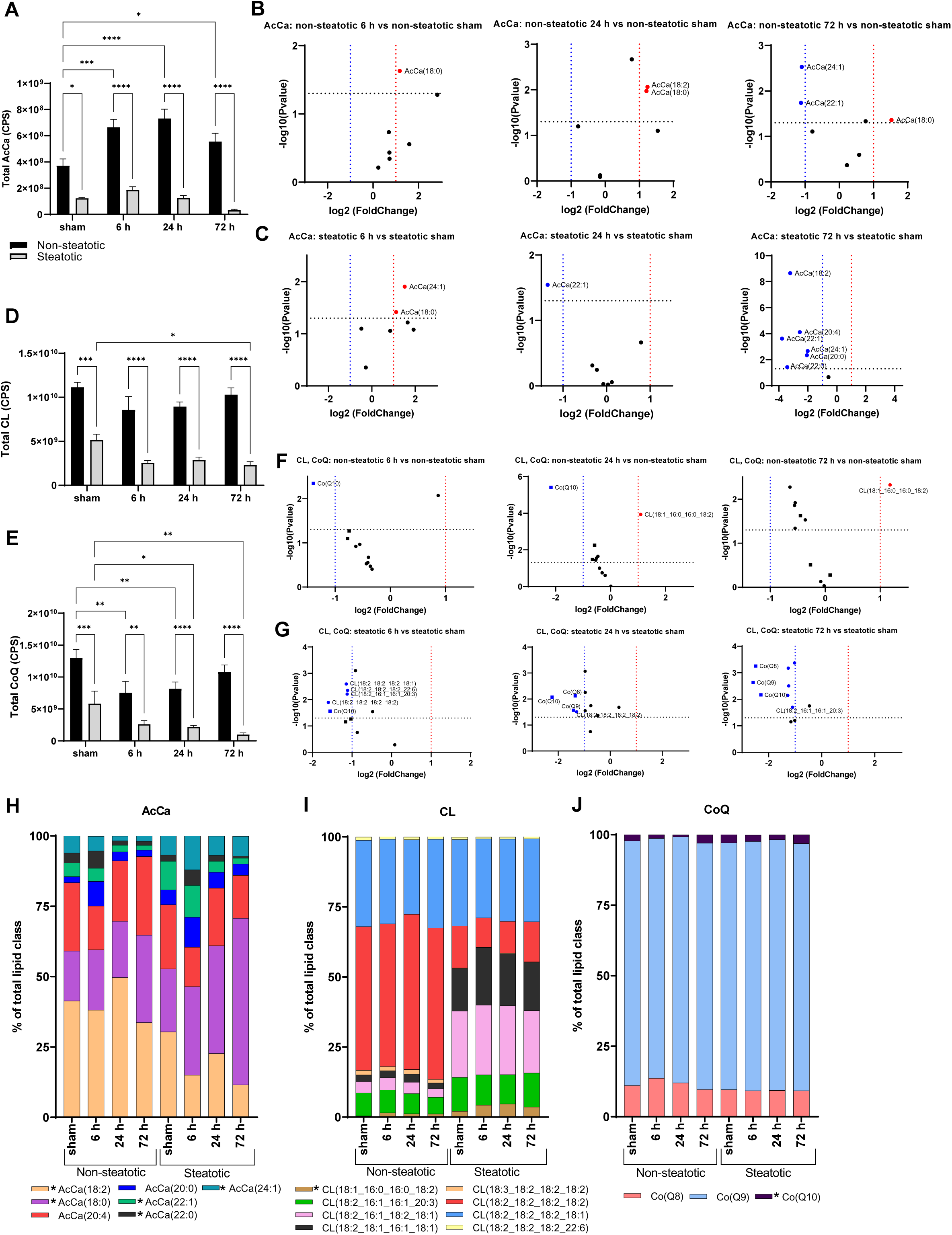
Analysis of mitochondrial lipids. **A**. Total acylcarnitines (AcCa) following sham or IR surgery. **B**. Volcano plot of specific AcCa species comparing non-steatotic 6 h, 24 h, and 72 h reperfusion to sham. **C**. Volcano plot of specific AcCa species comparing steatotic 6 h, 24 h, and 72 h reperfusion to sham. **D**. Total cardiolipin (CL) following sham or IR surgery. **E**. Total coenzyme Q (CoQ) following sham or IR surgery. **F**. Volcano plot of specific CL and CoQ species comparing non-steatotic 6 h, 24 h, and 72 h to CL and CoQ to sham. **G**. Volcano plot of specific CL and CoQ lipid species comparing steatotic 6 h, 24 h, and 72 h to sham. CoQ, squares, CL, circles. Specific AcCa [**H**], CL [**I**], CoQ [**J**] species represented as a percent of total lipid class following sham or IR surgery. *in legend indicates significant difference from respective diet sham I. Values are mean ± SEM. n = 4 per sham group, 8-10 per IR surgery group. Sham indicates sham surgery. 6 h, 24 h, 72 h indicates hours of reperfusion following IR surgery. *p<0.05, **p<0.01, p***<0.001, p****<0.0001. For volcano plots, lipids that are significantly increased relative to sham are represented in red, and those significantly decreased relative to sham are represented in blue. Black bars are chow fed mice and represent non-steatotic liver. Gray bars are 42% HF fed mice and represent steatotic liver.

Cardiolipin (CL) is an abundant component of mitochondrial membranes and is exclusively localized in this compartment. In non-steatotic liver, total CL did not change with IRI, but CL(18:1_16:0_16:0_18:2) was significantly increased at 24 h and 72 h compared to non-steatotic shams (Figure 6A, B). Coenzyme Q (CoQ) transports electrons in the electron transport chain. In non-steatotic liver, total CoQ decreased with IRI at 6 h and 24 h compared to non-steatotic sham (Figure 6D, E). In steatotic liver, there was a significant decrease in total CLs and total CoQ with IRI compared to steatotic sham (Figure 6D, E). Additionally, in steatotic liver, multiple CLs were significantly decreased at 6 h and 72 h, and all CoQs were decreased at 24 h and 72 h compared to steatotic sham (Figure 6F, G). Compared to steatotic liver, total CLs and total CoQs were higher in non-steatotic liver in sham and all reperfusion time points following IRI (Figure 6D, E).

We next looked at AcCa, CL, and CoQ species as a percent of total AcCa, CL, and CoQ, respectively (Figure 6H, I, J). In non-steatotic liver, the percentage of four AcCas changed with IRI relative to non-steatotic sham. In steatotic liver, the percentage of AcCa(18:0) significantly increased and AcCa(18:2) and AcCa(22:1) significantly decreased with IRI (Figure 6H). The composition of CLs in non-steatotic and steatotic liver was notably different at baseline. With IRI, the percentage of CL(18:1_16:0_16:0_18:2) increased in both non-steatotic and steatotic liver (Figure 6I). The proportion of each CoQ was similar between non-steatotic and steatotic liver. In both non-steatotic and steatotic liver, there was a significant decrease in the proportion of CoQ10 with IRI, but returned to baseline at 72 h reperfusion (Figure 6J). Collectively, these data indicate that IRI leads to a decrease in a majority of mitochondria-associated lipids in steatotic liver, but a similar decrease was not observed in non-steatotic liver.

### Comparison of sphingolipids

Sphingolipids play a variety of important roles in regulating intracellular signaling cascade and membrane dynamics and can also be classified into subtypes, including ceramides and hexosylceramides (Figure 7). Total ceramide content did not change with IRI at any time point in both non-steatotic and steatotic liver (Figure 7A). In non-steatotic liver there was a trend towards increased total HexCer content at 24 h and 72 h, but this did not reach statistical significance (p=0.37 and p=0.11, respectively) (Figure 7B). In steatotic liver, there was a significant decrease in total HexCer at 24 h post reperfusion compared to steatotic sham (Figure 7B). We did not detect any significant difference in total ceramide content between non-steatotic and steatotic liver in sham operated animals or at any reperfusion time point (Figure 7A). Non-steatotic liver had a significantly higher total HexCer content than steatotic liver at 24 h and 72 h post reperfusion (Figure 7B).

**Figure 7.**
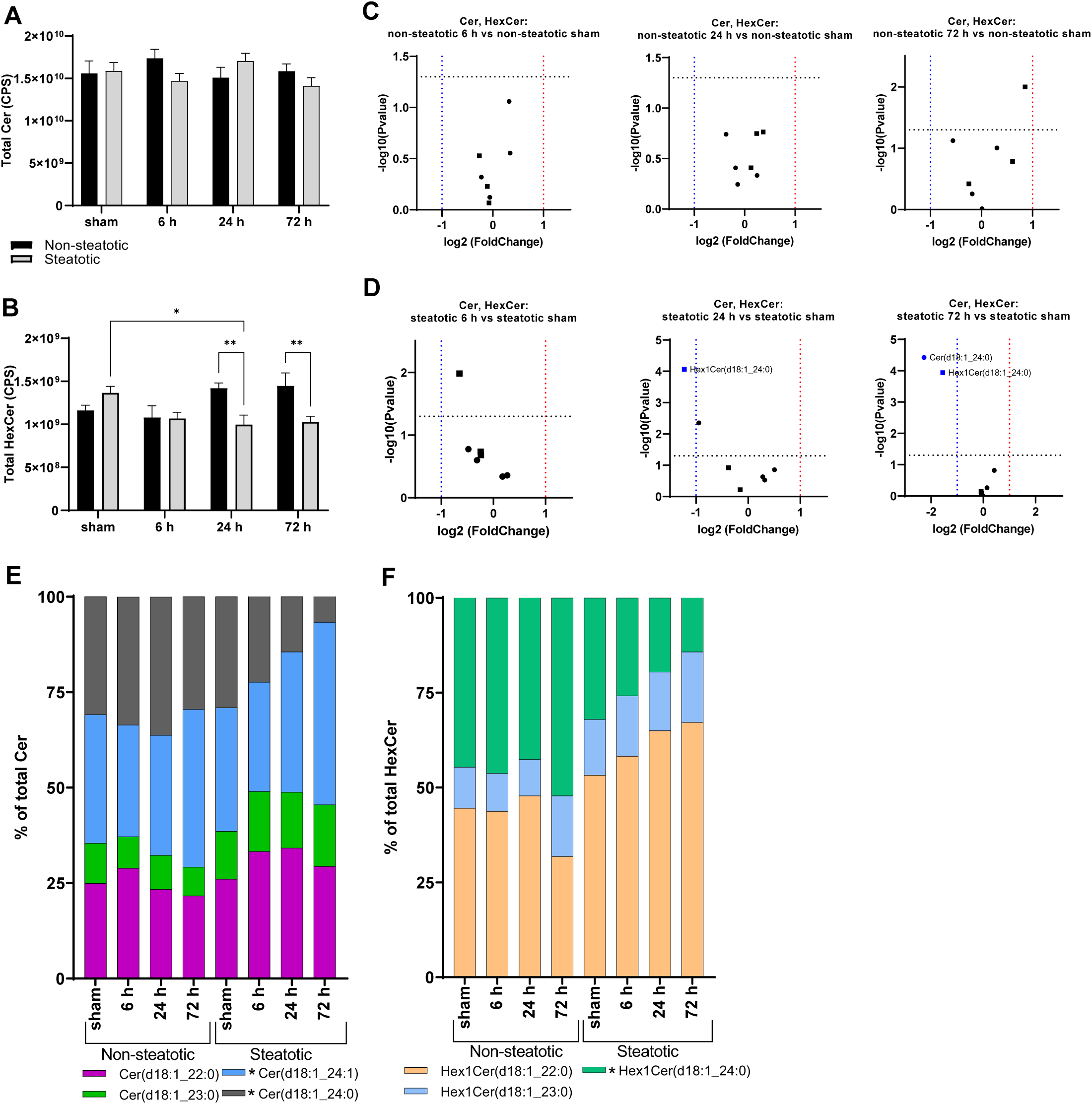
Analysis of sphingolipids. **A**. Total ceramide (Cer) following sham or IR surgery. **B**. Total hexosylceramide (HexCer) following sham or IR surgery. **C**. Volcano plot of specific Cer (circles) and HexCer (squares) species comparing non-steatotic 6 h, 24 h, and 72 h to sham. **D**. Volcano plot of specific Cer (circles) and HexCer (squares) species comparing steatotic 6 h, 24 h, and 72 h to sham. Specific Cer [**E**] and HexCer [**F**] species represented as a percent of total Cers and HexCer, respectively, following sham or IR surgery. *in legend indicates significant difference from respective diet sham. Values are mean ± SEM. n = 4 per sham group, 8-10 per IR surgery group. Sham indicates sham surgery. 6 h, 24 h, 72 h indicates hours of reperfusion following IR surgery. *p<0.05, **p<0.01, p***<0.001, p****<0.0001. For volcano plots, lipids that are significantly increased relative to sham are represented in red, and those significantly decreased relative to sham are represented in blue. Black bars are chow fed mice and represent non-steatotic liver. Gray bars are 42% HF fed mice and represent steatotic liver.

We then looked at individual ceramide species. In non-steatotic liver, none of the specific ceramide species exhibited a significant increase or decrease relative to non-steatotic sham following IRI at any reperfusion time point (Figure 7C). In steatotic liver, Cer(d18:1_24:0) and HexCer(18:1_24:0) were significantly decreased compared to steatotic sham (Figure 7D).

We then looked at each Cer and HexCer species as a percent of the total Cer and HexCer, respectively. In non-steatotic liver, there were no significant changes in the proportion of Cer or HexCer with IRI.

In steatotic liver, Cer(d18:1_24:0) and Hex1Cer(d18:1_24:0) decreased with IR injury, while Cer(d18:1_24:1) increased when compared to steatotic sham (Figure 7E, F). Together, these data suggest that sphingolipids were not significantly altered with IRI in non-steatotic liver while changes in steatotic liver were most notable for decreases in Hex1Cer(18:1_24:0) and Cer(d18:1_24:0).

### Correlation between lipid species and plasma alanine transaminase

As plasma ALT is the most common marker of liver injury, we next looked for correlations between ALT values and specific lipid species (Figure 8). Pearson correlation coefficients were calculated using all individual ALT and CPS values at each reperfusion time point. In non-steatotic liver, at 6 h reperfusion, ALT was significantly negatively correlated with LPI, LPC, PI, PC, PE, PS, PG, HexCer, CL, and CoQ. ALT was not significantly positively correlated with any lipid class measured at 6 h and 72 h. At 24 h, ALT was significantly negatively correlated with LPE and PE and positively correlated with AcCa (Figure 8A). In steatotic liver, at 6 h reperfusion, total LPI, LPC, PI, PG, HexCer, CL, and CoQ were significantly negatively correlated with plasma ALT. At 24 h reperfusion, none of the lipid classes were significantly associated with plasma ALT. At 72 reperfusion, only total CoQ was significantly negatively correlated with plasma ALT.

**Figure 8.**
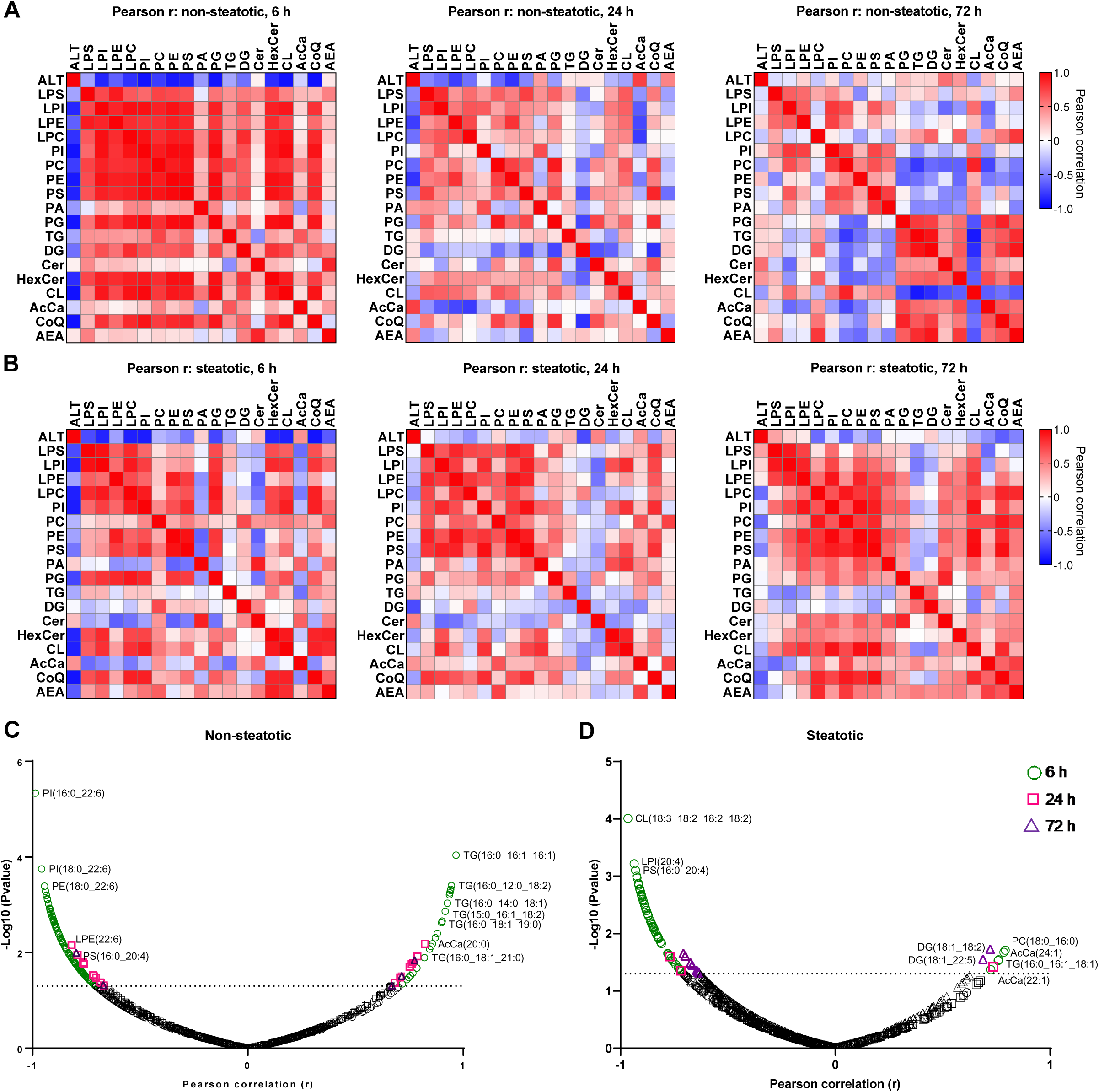
Pearson correlation matrix. **A**. Correlation between plasma ALT and lipid class in non-steatotic diet fed mice at 6 h, 24 h, and 72 h post reperfusion. **B**. Correlation between plasma ALT and lipid class in steatotic diet fed mice at 6 h, 24 h, and 72 h post reperfusion. Red indicates positive correlation. Blue indicates negative correlation. **C**. Pearson correlation between plasma ALT and specific lipid species in non-steatotic diet fed mice at all reperfusion time points. **D**. Pearson correlation between plasma ALT and specific lipid species in steatotic diet fed mice at all reperfusion time points. Dashed line on volcano plots indicate threshold for significant p-value (-Log10(pvalue)>1.3) plotted on y axis. Pearson correlation coefficient plotted on x axis.

As noted above, we often detected significant changes in individual lipid species even when no changes were detected from the total class after IRI. Thus, we examined each individual lipid species for correlation with plasma ALT in both non-steatotic and steatotic diet fed mice (Figure 8C, D). In non-steatotic liver, 138 individual lipid species were significantly correlated with plasma ALT at 6 h, 24 lipid species at 24 h, and five lipid species at 72 h. In steatotic liver, 64 lipid species were significantly correlated with plasma ALT at 6 h, three at 24 h, and eleven at 72 h reperfusion (Figure 8C, Supplemental Table 2).

We then examined how the correlated lipid species changed with IRI. In non-steatotic liver, at 6 h reperfusion, of the lipid species positively correlated with plasma ALT, two TG species (TG(16:0_12:0_18:2) and TG(16:1_14:0-18:2)) were significantly increased and one PI species (PI(18:0_22:5)) was significantly decreased relative to non-steatotic sham. Of the lipid species negatively correlated with plasma ALT at 6 h, four were significantly decreased and one was significantly increased (DG(18:3_18:2)) relative to non-steatotic sham. In non-steatotic liver, of the lipids positively correlated with plasma ALT at 24 h, AcCa(18:2) was significantly increased relative to non-steatotic sham and none were significantly decreased. Of the lipids negatively correlated with plasma ALT at 24 h, only three PEs were significantly decreased relative to non-steatotic sham and none were significantly increased. At 72 h reperfusion, while five lipid species were either negatively or positively correlated with plasma ALT, none of these lipids had a corresponding change relative to sham in non-steatotic liver.

In steatotic liver, of the lipid species that were positively correlated with plasma ALT, only AcCa(24:1) was significantly increased and none were significantly decreased at 6 h. In contrast, of the lipids that were negatively correlated with plasma ALT at 6 h, many were significantly decreased relative to steatotic sham. At 24 h reperfusion, 84 total lipids were significantly decreased following IRI in steatotic liver. However, only two of these lipids were significantly correlated with plasma ALT. Only AcCa(22:1) was positively correlated with plasma ALT in steatotic liver. This specific lipid was also significantly decreased in steatotic liver at 24 h. Of the lipids negatively correlated with plasma ALT in steatotic liver at 24 h, only PC(16:0_20:5) was also significantly decreased relative to steatotic sham. Of the lipid species negatively correlated with plasma ALT at 72 h, all except one (TG(20:1_18:2_22:6)) were significantly decreased at 72 h reperfusion compared to steatotic sham. Interestingly, of the 186 lipids that were significantly decreased following IRI in steatotic liver, only eight were correlated with plasma ALT.

Collectively, these data indicate that while many lipid species fluctuated following IRI, a relatively small percentage were correlated with plasma ALT. Of these correlations, a general pattern was noted whereby phospholipids were negatively correlated with plasma ALT and in general tended to decrease following IRI. This finding was more pronounced in steatotic liver and persisted at the 72 h reperfusion time point.

## Discussion

Ischemia reperfusion injury is largely unavoidable in most liver-related surgeries, and the presence of steatosis exacerbates injury. There are no specific biomarkers used for diagnosing or prognosticating hepatic IRI, and there are no pharmacological therapies available to prevent or treat IRI. While it is generally thought that accumulation of toxic lipid species is linked to hepatic inflammation and fibrosis (21, 22), how specific lipids impact IRI is less well understood. In this study, we used an unbiased, untargeted approach to systematically evaluate changes in lipid composition following IRI in both non-steatotic and steatotic liver in mice.

Broad analysis indicated that steatotic and non-steatotic liver have distinct changes to intrahepatic lipid profiles following IRI. This highlights the need to perform studies in both non-steatotic and steatotic liver, as a positive response to intervention in one may not be applicable to the other. Interestingly, even when content of a given total lipid class did not change significantly with IRI, we noted fluctuations in specific lipids within that class. These changes in composition may reflect fatty acid availability or substrate preferences. This will require further investigation as previous studies have indicated that specific fatty acids may influence IRI outcomes (23, 24). Additionally, we noted that there were very few lipids that increased relative to sham following IRI in either non-steatotic or steatotic liver. Of note, PA(18:0_20:4), the only PA detected in our study, was increased at all reperfusion time points in steatotic liver, but unchanged in all time points in non-steatotic liver. PA has mitogenic effects and may play a role in liver regeneration (11, 25–29). The increase in PA in steatotic liver may be a reflection of increased liver injury, and thus a need to initiate regeneration following IRI.

In addition to PA, TG was one of the few lipid species that increased following IRI. Specifically, we noted significant TG accumulation in non-steatotic liver following IRI. This is in line with previous findings of transient steatosis following IRI (11) and partial hepatectomy (30). Of note, we did not see a similar increase in TG in steatotic liver. The differences in TG accumulation may be secondary to differences in fatty acid oxidation, lipophagy, lipolysis, TG synthesis or a combination of these factors. Future mechanistic studies will be needed to address mechanisms and significance of this finding.

With the exception of PA, total phospholipids and glycerophospholipids, specifically total PC, PE, PI, PS, LPC, and LPE, all decreased with IRI in steatotic liver following IRI. Furthermore, we found multiple PC and PE lipid species to be negatively correlated with plasma ALT, suggesting that decreased phospholipid content is associated with increased liver injury. Phosphatidylcholine has an essential role in the assembly of cell membranes, lipoproteins, lipid droplets and bile synthesis (31). Aberrant phosphatidylcholine metabolism has been linked to NAFLD and liver failure (32–34), cardiovascular disease, myocardial ischemia reperfusion injury (35, 36), and Alzheimer’s disease (37, 38). Furthermore, liver regeneration following partial hepatectomy is influenced by the PC:PE ratio (39) and similar to our findings, in the murine acetaminophen model, almost all PC species decreased following acetaminophen treatment (40). A recent multiomics study by Hall Z, et al (41) suggest that hepatocyte proliferation is associated with increased de novo synthesis of PC. The role of phospholipid metabolism has not been specifically investigated in hepatic IRI, but PC administration has been shown to decrease intestinal IRI (42) and brain IRI (43). While Zazueta et al found CDP-choline to ameliorate IRI in non-steatotic liver, they did not evaluate effects in steatotic liver and did not measure lipid levels (44). Given the important role of PC as the predominant phospholipid in cell membranes, the relative decrease in PCs following IRI compared to non-steatotic livers may contribute to slower resolution of and recovery from IRI in steatotic liver. Thus, future studies will need to evaluate PC content as prognostic markers and the role of PC supplementation in hepatic IRI.

Interestingly, we found that AcCa was positively correlated with plasma ALT in both non-steatotic and steatotic liver. Changes in AcCa likely result from alterations in fatty acid oxidation seen in ischemia reperfusion injury (45, 46). To our knowledge, there have not been previous studies evaluating a relationship between AcCa and hepatic IRI. Previous work has reported beneficial effects of L-carnitine in rat hepatic IRI (47). Liepinsh et al previously found decreasing acylcarnitine content was associated with decreased myocardial IRI (48). However, future studies evaluating the effects of AcCa on hepatic IRI are needed.

In addition to our novel findings, our study also reports changes that are consistent with previous studies. For example, we found cardiolipin, a phospholipid exclusive to the mitochondria, to be significantly decreased in steatotic liver following IRI and negatively correlated with plasma ALT. Multiple groups in cardiac (49) and hepatic (12) IRI have reported similar findings suggestive of mitochondrial dysfunction secondary to IRI. We also noted decreased CoQ following IRI and previous studies have noted alleviation of IRI with administration of CoQ10 (50–52). Of note, previous studies (10, 53–55) have noted changes in ceramide content following IRI, but we did not detect such alterations. This may be due to differences in study design and methodology.

There are several limitations to our study. Importantly, this study is descriptive in nature. While not intended to define the mechanisms by which alterations in lipid composition occur, it lays the foundation for design of future mechanistic studies. Further studies including transcriptomic analysis are needed to fully evaluate how altered lipidomics may contribute to IRI and how such changes may be exploited for therapeutic potential. Second, we only utilized one murine model NAFLD. However, our model is most relevant as genetic models of obesity and the methionine choline deficient diet are less physiologically applicable. Additionally, the NAFLD produced by the 42% HF diet creates a reliable, reproducible model of NAFLD with steatosis and inflammation without significant fibrosis. Third, female mice were not included in this study, and there may be sex-specific responses to IRI. Future targeted lipidomic studies should include both male and female mice to explore such differences. Finally, in this study, we did not perform comprehensive plasma lipidomics. However, our primary aim was to characterize hepatic lipidomic changes as this is the site of primary injury. Plasma lipidomics would reflect more systemic changes or changes to other organs such as adipose tissue following IRI. In the future, it would be important to further delineate changes in plasma lipids as this would allow for study of systemic effects of IRI and help identify novel non-invasive biomarkers.

It is thought that steatotic livers are more susceptible to IRI due to underlying mitochondrial dysfunction, ER stress, and microcirculatory impairment that exaggerates liver injury and cell death following IRI. Hence, most studies have focused on interventions that would target these pathways. However, dysregulated lipid metabolism and alterations in lipid composition have been shown to contribute to various disease states. In our present study, utilizing unbiased, comprehensive lipidomic analysis, we have illustrated that there are distinct and dynamic changes to lipid profiles following IRI in non-steatotic and steatotic liver. Specifically targeting lipid metabolism represents a novel therapeutic approach. Our findings expand our knowledge of the lipidomic changes that occur and provide valuable insight regarding biomarker identification and therapeutic strategies.

## Data availability statement

All data are contained within the manuscript. Data requests can be made to Brian Finck, Washington University School of Medicine, at.

## Abbreviations

42% HF: 42% high fat diet Envigo TD.88137
AcCa: acylcarnitine
ANOVA: analysis of variance
AU: arbitrary unit
ALT: alanine aminotransferase
Chow: standard chow diet, PicoLab Rodent Diets 20 5053
CL: cardiolipin
CPS: counts per second
DG: diacylglycerol
FC: fold change
Hex1Cer: hexosylceramides
Il: interleukin
IR: ischemia reperfusion
IRI: ischemia reperfusion injury
TNF: tumor necrosis factor
KEGG: Kyoto encyclopedia of genes and genomes
LC/MS: liquid chromatography mass spectrometry
NAFLD: nonalcoholic fatty liver disease
PC: phosphatidylcholine
PCA: principal component analysis
PE: phosphatidylethanolamine
PI: phosphatidylinositol
PS: phosphatidylserine
LPC: lysophosphatidylcholine
LPE: lysophosphatidylethanolamine
LPI: lysophosphatidylinositol
SEM: standard error of the mean
TG: triglyceride

