## supplemental figures for "Dynamic changes in the mouse hepatic lipidome following warm ischemia reperfusion injury"

Supplemental Figure 1. Body weight and liver histology of mice fed a chow or 42% HF diet.

Supplemental Figure 2. PA content following sham or IR surgery.

Supplemental Figure 3. Complete legends corresponding to Figures 3G, 4M, 4N, 4O.

*indicates significant difference from sham

Supplemental Table 1. Lipid species significantly increased or decreased relative to sham.

Supplemental Table 2. Lipid species with significant positive or negative correlation with plasma ALT.

Supplemental Figure 1.

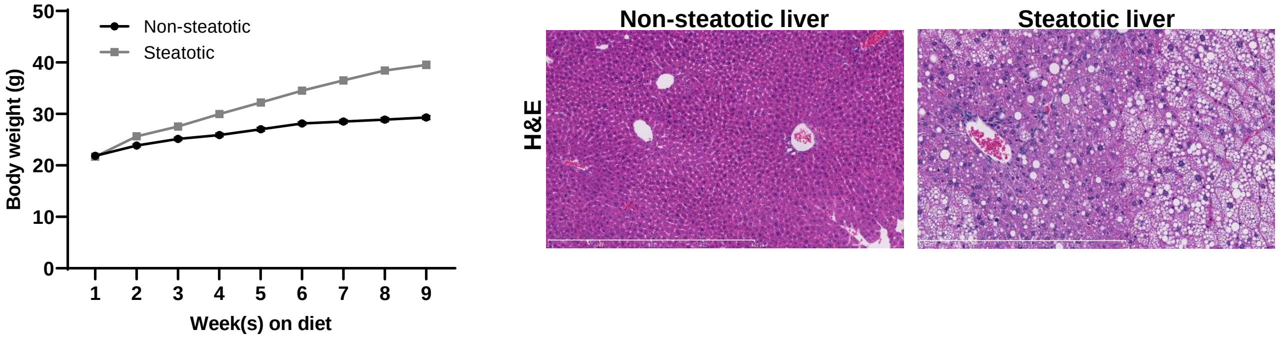

Supplemental Figure 2.

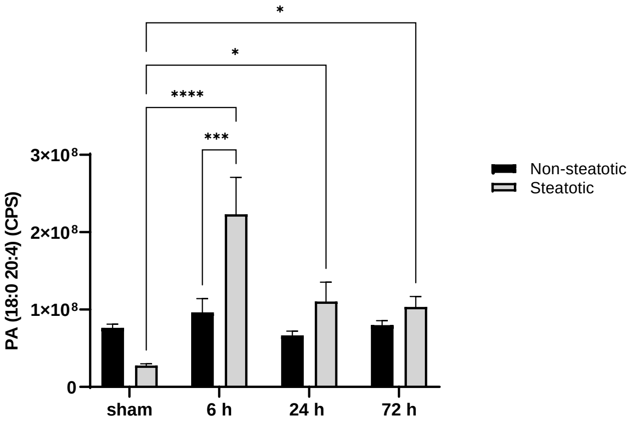

Supplemental Figure 3.

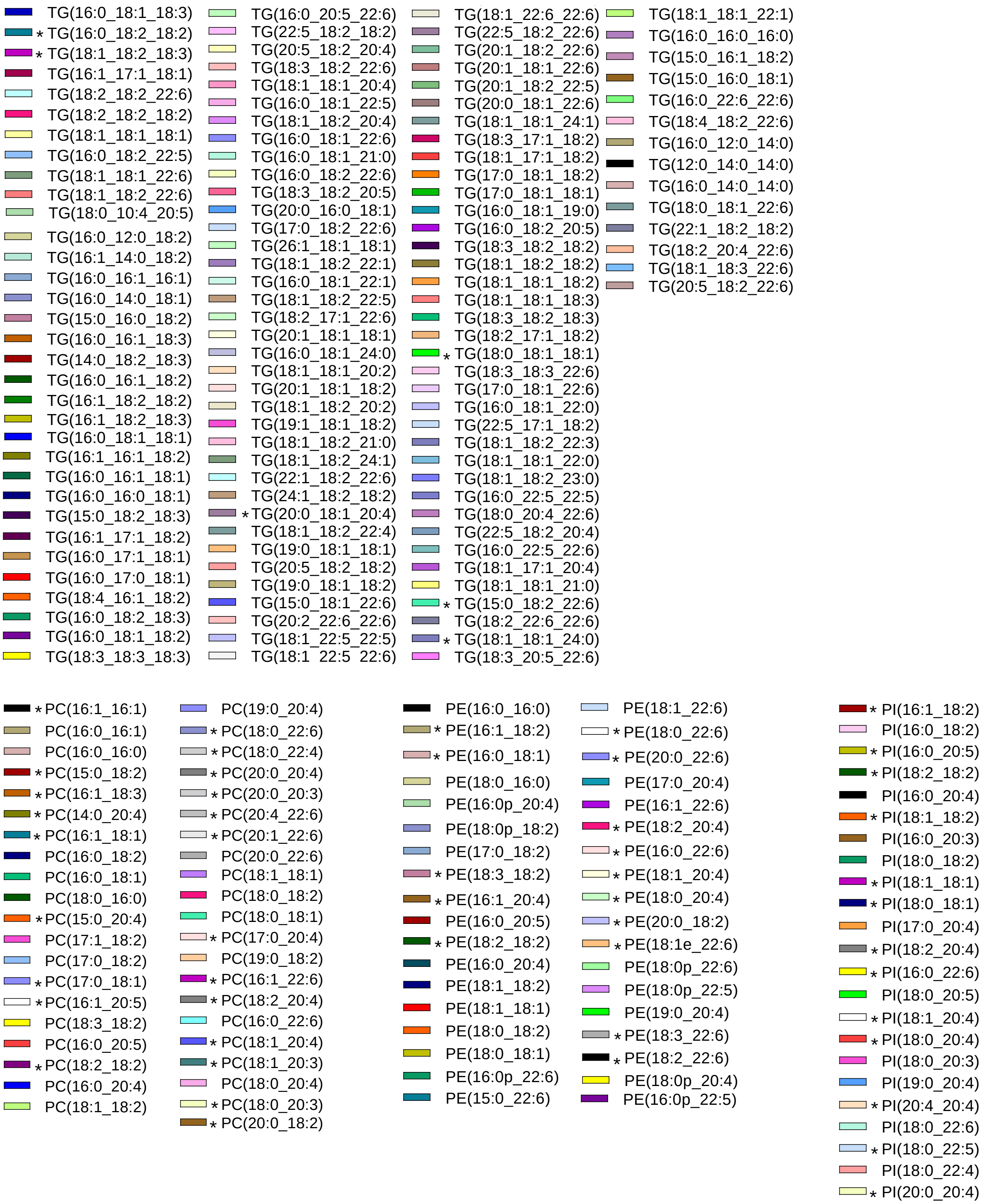
